# Cleavage specificity of *E. coli* YicC endoribonuclease

**DOI:** 10.64898/2026.03.25.714237

**Authors:** Sarah A. Barnes, Michael B. Lazarus, David H. Bechhofer

## Abstract

*Escherichia coli* YicC enzyme is the founding member of a family of endoribonucleases that is encoded in virtually all bacterial species. Previous structural studies revealed that this ribonuclease binds RNA by a novel mechanism in which the hexameric apoprotein presents an open channel that undergoes a large rotation upon RNA binding and clamps down on the RNA. The current study follows up on these findings by examining the cleavage of various oligonucleotide substrates designed to probe recognition elements required for YicC binding and cleavage. A 26-nucleotide RNA oligomer (oligo), with a K_D_ in the low micromolar range, was the standard to which numerous oligos with altered sequence were compared. *In vitro* RNase assays and fluorescence anisotropy binding measurements indicated that the preferred substrates for YicC were relatively small RNAs that contain some secondary structure. Larger RNAs or highly structured RNAs were less-than-optimal substrates. Similarly, RyhB RNA, a ∼90-nucleotide, iron-responsive RNA of *E. coli*, which has been described as a target of YicC binding and/or cleavage, was a poor YicC substrate in our assays. These results suggest that the native substrates for YicC-family members are very small RNAs or RNA fragments derived from larger RNAs.

## INTRODUCTION

Despite decades of research on ribonucleases in *Escherichia coli* and *Bacillus subtilis*, new ribonucleases apparently remain to be found. Our group recently used an extract of a “quadruple mutant” of *B. subtilis*, in which genes for all four known exoribonucleases were knocked out, to discover a new ribonuclease, YloC (Ingle et al. 2022). We and others (Huang et al. 2023) found that YloC is active as a hexameric endoribonuclease. A similar observation was made for the homologous *Clostridiodies difficile* CD25890 protein (Martins et al. 2021). YloC is a member of the “YicC family” of proteins, named after the prototypical family member in *E. coli*; YicC homologues are present in virtually all bacterial species. YloC and YicC catalyze the identical endonuclease reaction on a 26-mer oligonucleotide (oligo) that carries a 5’ end IR-fluorescent label (Wu et al. 2024). An X-ray crystal structure for the apo-YicC protein and a cryo-EM structure for a YicC-RNA complex, using the same RNA oligo (unlabeled) that was used for endonuclease assays, revealed that the unbound hexamer exists in a conformation akin to an open clamshell, while the RNA-bound structure exists in a closed clamshell-like structure (Wu et al. 2024). This profound change in conformation upon RNA binding is unique among ribonucleases.

Using mass spectroscopy analysis to identify YicC cleavage products, it was determined that YicC cleaves the 26-nt RNA oligo endonucleolytically at two sites: a 5’-proximal site 1, between nts 3 and 4, and a downstream site 2, between nts 14 and 15 (Fig. 1A, 1B) (Wu et al. 2024). The 5’-proximal cleavage occurs in the upstream side of a hairpin structure, consisting of a 4-base-pair stem and 7-nucleotide (nt) loop, as seen in the cryo-EM structure. Cleavage at this site appears to be preferred. The downstream cleavage site occurs in a predicted hairpin structure, consisting of a putative 4-base-pair stem and a 3-nt loop. We proposed that the RNA oligo can be bound by YicC in one of two conformations, presenting either site 1 to the enzyme catalytic site, which is the conformation determined by cryo-EM, or presenting site 2 to the catalytic site. The 23-nt fragment remaining after site 1 cleavage was proposed to be also a substrate for site 2 cleavage by YicC. In the current investigation, we performed enzymatic assays of 5’-end labeled or 3’-end labeled oligos with different sequences, to probe the requirements for YicC binding and endonuclease activity.

**Fig. 1.**
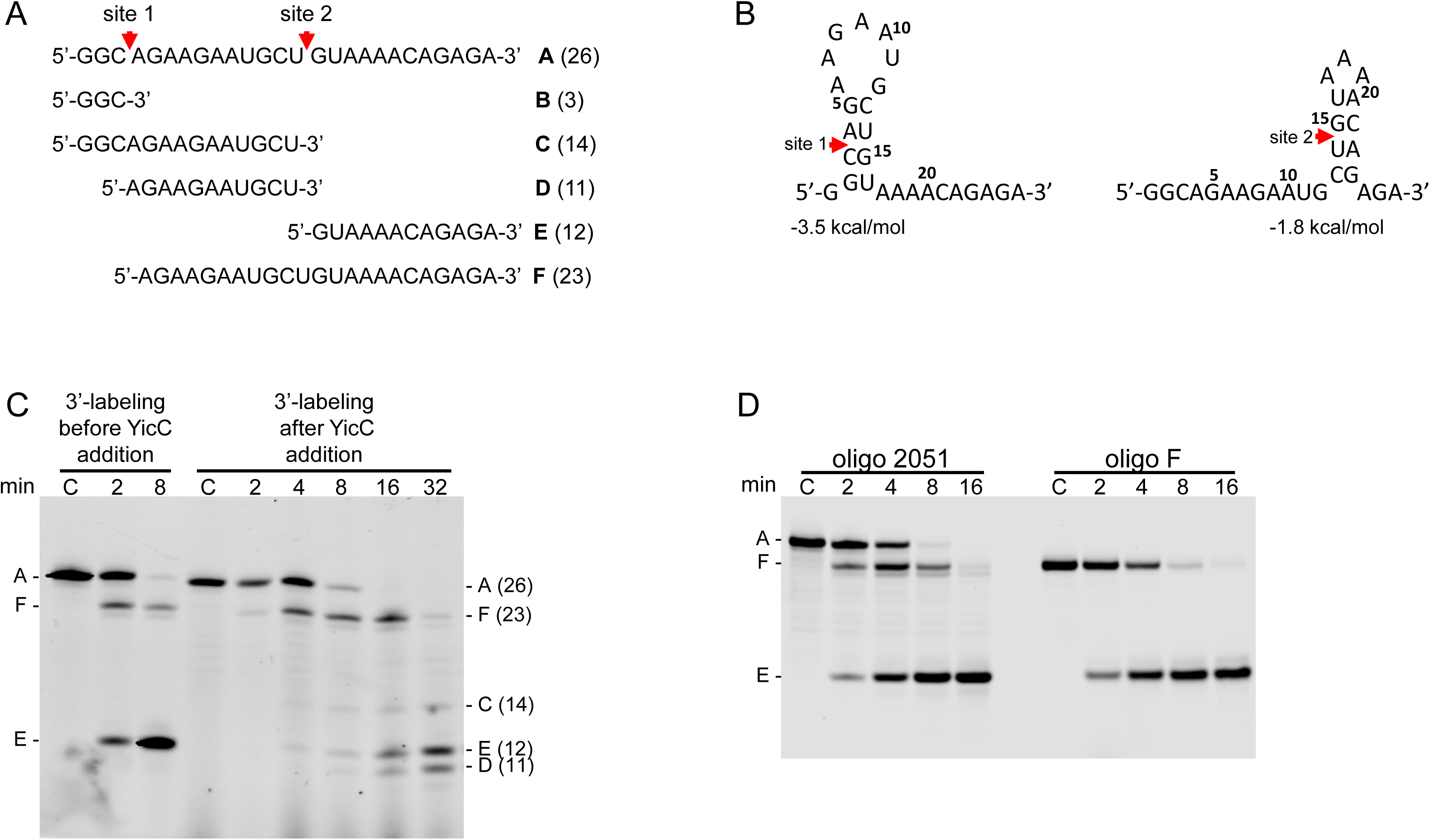
(A) Sequence of oligo 2051, showing two YicC cleavage sites and all possible fragments that result from cleavage at sites 1 and 2; number of nucleotides in each fragment indicated in parentheses. (B) Predicted alternate structures with 5’-proximal or 3’-proximal hairpins, with cleavage sites 1 and 2 indicated, and stability in kcal/mol as predicted by Mfold (Markham and Zuker 2008). (C) Denaturing gel electrophoresis (20% polyacrylamide) showing 3’-end labeled YicC cleavage products for oligo 2051, labeled before (left) and after (right) incubation for indicated times with YicC. Migration of cleavage products indicated on the left and right, with predicted size of each fragment shown in parentheses. Time in min indicated above each lane. Control lanes (C) contained oligo 2051 with no YicC added. (D) Cleavage of labeled F fragment.

## RESULTS

### 3’-end labeling to observe cleavage products

In our initial study of YicC cleavage activity, we used a 5’-end IR-fluorescent labeled, 26-nt RNA oligo to observe, by gel electrophoresis, the cleavage products of endonuclease activity. This standard oligo was designated “2051” and its sequence is shown in Fig. 1A. As this oligo was labeled at the 5’ end, we could observe only the cleavage products that retained the 5’ end (i.e., fragments B and C in Fig. 1A). We could not observe fragments that did not contain the 5’ end but that were identified by mass spectroscopy (Fig. 1A, fragments D, E, and F). To confirm that all expected fragments were generated in the YicC cleavage reaction, we used a technique to label RNA 3’ ends using fluorescein 5-thiosemicarbazide (FTSC) (Zearfoss and Ryder 2012). Thus, oligo 2051 was labeled at its 3’ end either before incubation with YicC or at increasing times after addition of YicC. For the RNA labeled before the YicC reaction, the results (Fig. 1C, left) showed the full-length oligo and two expected fragments, F and E, that retain the 3’ end after cleavage at sites 1 and 2, respectively. For the RNA labeled after addition of YicC, we observed (Fig. 1C, right) the full-length oligo and four other cleavage products that had been labeled at their 3’ ends. These RNA fragments migrated as expected for their predicted sizes (fragments F, C, E, and D). (We could not see the small 3-nt product, fragment B, as this product cannot be recovered after the ethanol precipitations that are part of the labeling protocol.) These results confirm, by direct observation, the YicC-catalyzed cleavage pattern for this oligo.

We hypothesized previously that oligo 2051 could adopt either of two alternate structures (Fig. 1B), which are mutually exclusive. The structure that contained the 5’-proximal stem-loop (nucleotides 2-16) was preferred, and after cleavage at site 1 the remaining RNA (fragment F) could form the structure with a stem-loop comprising nucleotides 13-23. The low abundance of fragment C (Fig. 1C) is consistent with this hypothesis, as it represents cleavage of the full-length RNA at site 2, which we do not expect to be a frequent event. The F fragment, which is produced by cleavage at site 1, increases in intensity early on but then decreases until almost none is present at the end of the time course. At the same time, fragments D and E increase in intensity with time of the reaction. This is consistent with the hypothesis that fragment F can be cleaved at site 2, generating fragments D and E. To show this directly, we used a 23-nt oligo that had the sequence of the F fragment only (oligo F) and contained a 5’ phosphoryl group (as would the F fragment that is generated by YicC cleavage of full-length oligo 2051). Oligo F was labeled at its 3’ end and incubated with YicC. The results (Fig. 1D) demonstrate that the F fragment is cleaved efficiently by YicC. Thus, YicC cleavage at site 2 is not dependent on initial binding to the full-length RNA with its 5’-proximal stem-loop structure.

### Effect of ribose 2’-modifications at the cleavage site 1

We showed previously that *B. subtilis* YloC is a divalent cation-dependent enzyme (Ingle et al. 2022). For metal-dependent enzymes, the 2’-OH position is not involved in the catalytic mechanism. To probe the catalytic mechanism of YicC, we designed RNA oligos that have the same sequence as oligo 2051 but have a 2’-deoxyribonucleotide at either nt 3 or nt 4, designated 3(2’-H) and 4(2’-H). As shown in Supplementary Fig. 1A and 1B, when either of these oligos was labeled at the 5’-end with 5-(iodoacetimado)fluorescein (5-IAF) or the 3’ end with FTSC, YicC was able to cleave at the same positions as for oligo 2051. This result supports the notion that *E.coli* YicC is also a metal-dependent enzyme, as has already been shown by Huang et al. (Huang et al. 2023).

On the other hand, when we tested oligos in which both nts 3 and 4 were either deoxyribonucleotides (designated 3,4 (2’-H)) or 2’O-methylated nucleotides (designated 3,4 (2’-OM)), cleavage at site 1 was abolished, while cleavage at site 2 was maintained (Supplementary Fig. 1C and 1D). Presumably, the absence of 2’-hydroxyl groups in nucleotides 3 and 4, which form part of the 5’-proximal stem sequence, affected structure to the extent that this was no longer recognized by YicC (Darre et al. 2016). These results support the hypothesis that secondary structure, rather than nucleotide sequence alone, is a necessary element for YicC substrates.

### Effect of stem-loop perturbations

The hypothesis that oligo 2051 can adopt either of two conformations, each of which is cleaved on the 5’ side of a stem-loop structure, predicted that it should be possible to observe substrates that are cleaved at a single site. Thus, we designed 26-nt oligos that have sequence changes from oligo 2051 that are predicted to disfavor formation of one or the other stem-loop structure. Oligo 3(A) was changed at position 3 from a C to an A residue (Fig. 2A). This change disrupts the predicted 5’-proximal stem-loop base pairing but not the predicted 3’-proximal stem-loop base-pairing. Indeed, oligo 3(A) was cleaved at site 2 but not at site 1 (Fig. 2B). Similarly, we tested oligo 22-24(GAC), which differs from oligo 2051 at nts 22-24. This change is predicted to disrupt the 3’-proximal stem base-pairing but not affect the 5’-proximal stem base-pairing. As shown in Fig. 2B, oligo 22-24(GAC) was cleaved at site 1 but barely at site 2. Oligo 4-6(GAC) had the AGA sequence of nts 4-6 of oligo 2051 replaced with a GAC sequence. This change was predicted to result in the formation of a 5’-proximal structure with a 4-bp stem and 4-nt loop, with the stem sequence beginning five nts from the 5’ end rather than 2 nts as for oligo 2051 (Fig. 2A). The cleavage pattern for oligo 4-6(GAC) was the same as for oligo 2051, except that the site 1 cleavage fragment was larger. Finally, we tested activity on an oligo that switched the 5’ and 3’ sections of oligo 2051, such that nts 13-26 of oligo 2051 were at the 5’end and nts 1-12 of 2051 were at the 3’ end. We designated this the “switched” oligo, whose predicted structure is shown in Fig. 2A. The RNase assay showed that switched oligo was cleaved closer to the 5’ end, likely between the second and third nucleotide of the upstream side of the stem structure. This is the same relative position as site 1 of oligo 2051 (cf. Fig. 1B), even though the nucleotide sequence is different. These results demonstrate that YicC cleavage is targeted specifically to double-stranded regions.

**Fig. 2.**
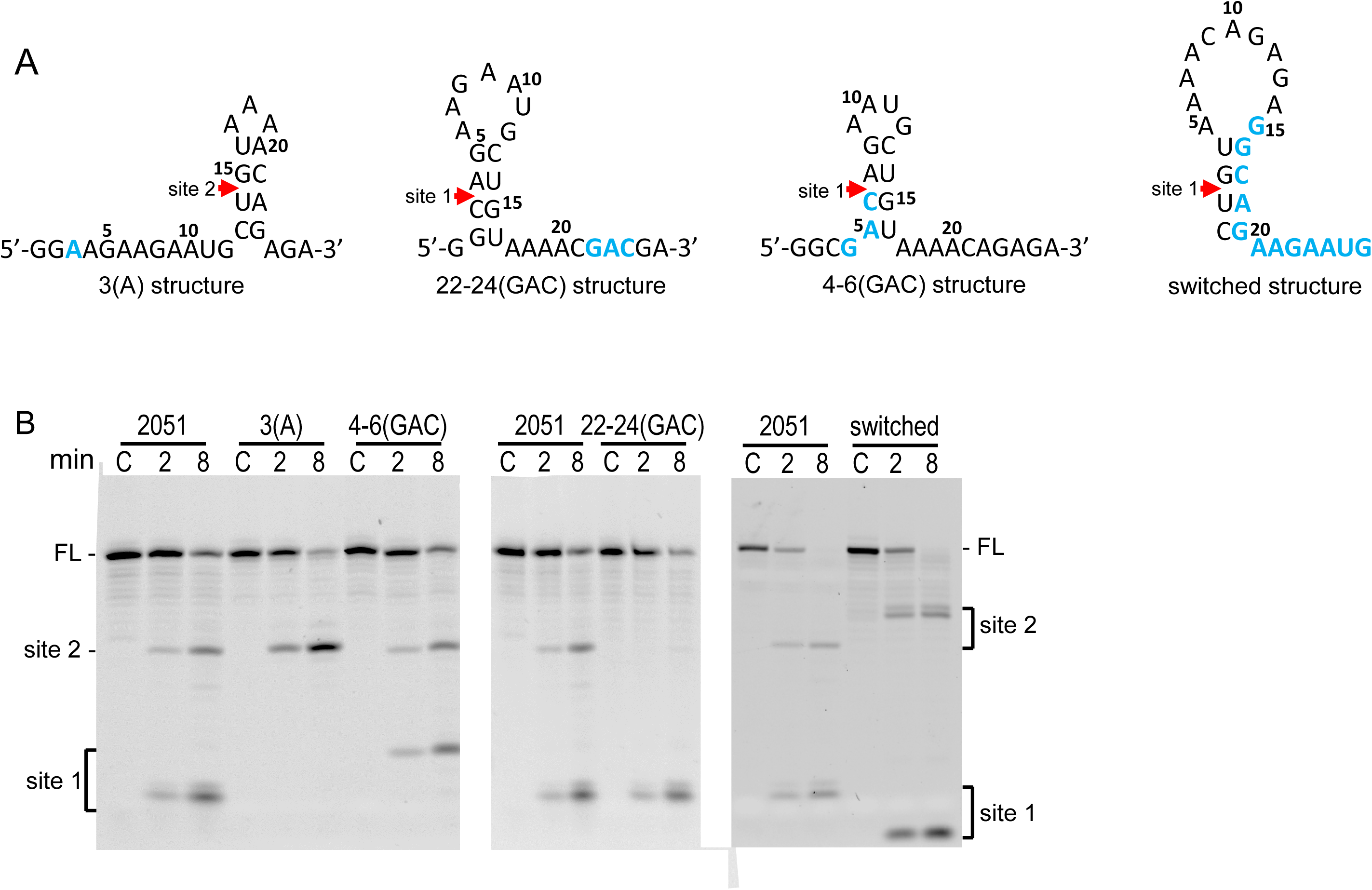
(A) Predicted secondary structure of mutated versions of oligo 2051. Shown are the most stable structures as predicted by Mfold. Nucleotide changes in oligos 3(A), 22-24(GAC), and 4-6(GAC), relative to oligo 2051, are shown in blue, with cleavage site 1 or 2 indicated. For the switched oligo, nucleotides 1-12 of oligo 2051 are shown in blue. (B) Denaturing gel electrophoresis (20% polyacrylamide) showing YicC cleavage products of 5’-end labeled oligos with sequence changes from oligo 2051. Migration of full-length (FL) oligo and products of cleavage at sites 1 and 2 are indicated at left and right. Time in min indicated above each lane. Control lanes (C) contained oligo with no YicC added.

### YicC activity on oligos with stem-loop structures of altered stability

Having demonstrated the requirement for a stem-loop structure, we probed whether increasing the predicted stability of the 5’-proximal stem-loop structure – by adding additional base pairs – would affect YicC activity. To simplify the analysis, we first created an oligo in which the final 10 nts were all A residues (Fig. 3A). (Although a sequence of consecutive A residues can form its own local helical structure whereas consecutive U residues do not form structures (Hashizume and Imahori 1967), we could not use oligos with U residues at the 3’ end. When we examined the YicC cleavage activity on an oligo with 10 U residues at the 3’ end, we observed multiple cleavage sites, likely because of alternative secondary structure formation, e.g., U-A and U-G base-pairing.) The presence of only A residues at the 3’ end was predicted to eliminate cleavage at site 2 but not alter secondary structure formation and cleavage at site 1. Thus, this oligo was designated “5’stem,” as it was predicted to be able to form only the 5’-proximal stem. Indeed, 3’-labeled oligo 5’stem was cleaved at site 1 only (Fig. 3B). There did appear to be somewhat less specificity in the exact cleavage site, relative to oligo 2051.

**Fig. 3.**
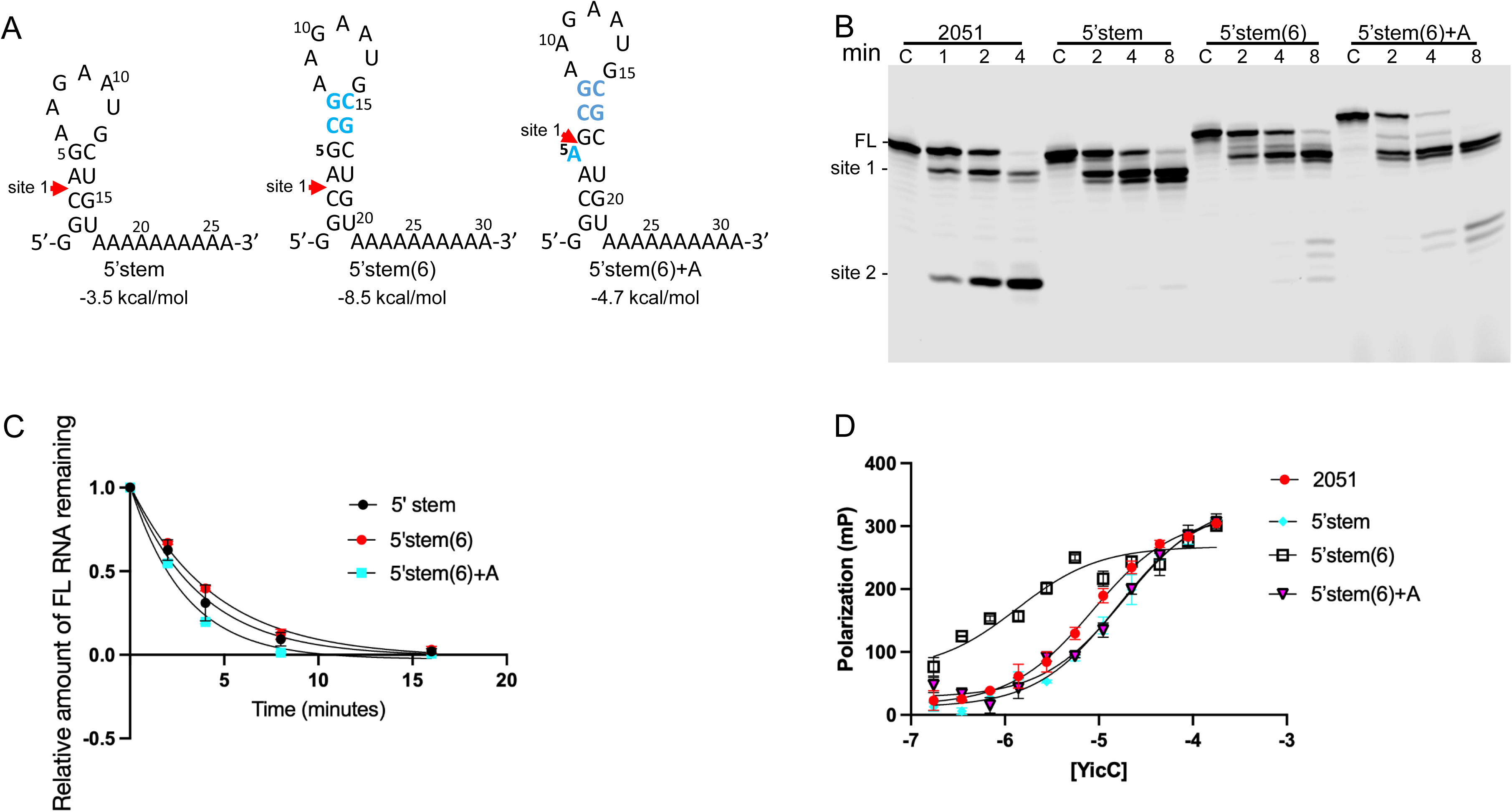
(A) Predicted secondary structures of oligos with 10 A residues at the 3’ end that precludes formation of the 3’-proximal cleavage site. Oligo 5’stem has the same upstream sequence as 2051; oligos 5’stem(6) and 5’stem(6)+A have additional nucleotides inserted, shown in blue. (B) YicC cleavage assay on 3’-labeled oligos; migration of full-length and cleavage products for oligo 2051 indicated on the left. Time in min indicated above each lane. Control lanes (C) contained oligo with no YicC added. (C) Decay rate of oligos 5’stem, 5’stem(6), and 5’stem(6)+A. Average of three experiments. (D) Fluorescence anisotropy of 3’-labeled oligos with wild-type YicC protein. Average of three experiments.

Next, four nucleotides were added that were predicted to form two extra base-pairs (for a total of six) in the 5’-proximal stem-loop structure, giving oligo 5’stem(6) (Fig. 3A). The predicted stability, by Mfold, of 5’stem(6) was five orders of magnitude stronger than for oligo 2051. The RNase assay on 5’stem(6) showed full-length RNA and a cleavage product whose migration was consistent with the additional four nucleotides. The cleavage rate was slightly slower than for the 5’stem oligo (Fig. 3C). Next, an extra A residue was inserted to introduce a predicted bulge in the 6-bp stem of 5’stem(6), giving oligo 5’stem(6)+A (Fig. 3A). The predicted stability of oligo 5’stem(6)+A was closer to that of oligo 2051. Surprisingly, oligo 5’stem(6)+A was cleaved at a faster rate than the other two oligos (Fig. 3C). To determine the approximate position of site 1 cleavage of these oligos, they were 5’-end labeled and the relative migration of the small, 5’-terminal fragment was observed (Supplementary Fig. 2). While the position of the cleavage site was the same for the 5’stem and 5’stem(6) oligos, which was determined previously to be 3 nts from the 5’ end (Wu et al. 2024), for the 5’stem(6)+A oligo the cleavage appeared to be 2-3 nucleotides further downstream. The fact that oligo 5’stem(6)+A, with the bulged stem-loop, was cleaved at a faster rate than the other two oligos in this set suggested that YicC prefers a non-perfectly paired stem as a substrate.

To determine whether changes in the rate of cleavage were due to differences in substrate binding or to cleavage activity, we performed fluorescence anisotropy measurements, using the 3’-end labeled oligos that were used in the cleavage assay but that had been purified by gel extraction. The binding affinity (K_D_) for oligo 5’stem was about two-fold higher than for oligo 2051 (Fig. 3D, Table 1). The K_D_ for the 5’stem(6) oligo, which was predicted to have six, rather than four, base-pairs in its stem-loop structure, was about six-fold lower than for oligo 5’stem. Nevertheless, oligo 5’stem(6) was cleaved at a slightly slower rate than the 5’stem oligo (Fig. 3C). In other words, the increased stability of the stem-loop structure in the 5’stem(6) oligo made for a stronger binding affinity, but, nevertheless, the cleavage rate for 5’stem(6) was the slowest of the three oligos. 5’stem(6)+A, in which the inserted A nucleotide was predicted to result in a bulge, had a similar K_D_ to that of the 5’stem oligo, but, nevertheless, was cleaved at the fastest rate.

**Table 1.** IC50 values for 3’-end labeled oligos. Absolute K_D_ values cannot be compared from one fluorescence anisotropy experiment to another because different batches of purified YicC protein were used. Thus, we show Kd values for the different oligos relative to the value for oligo 2051 in an individual experiment. These are the average of three measurements, except for the 5’+5 and 5’+10 oligos which were the average of six measurements. The actual K_D_ value for oligo 2051 was in the 5-15 μM range.

| <u>Oligo name</u> | <u>Fold-change from 2051 $K_D$ value</u> |
| --- | --- |
| 2051 | 1.00 |
| 5'stem | 1.83 |
| 5'stem(6) | 0.16 |
| 5'stem(6)+A | 2.17 |
| 5'+5 | 1.79 |
| 5'+10 | 1.24 |
| 3'+5 | 1.54 |
| 3'+10 | 3.89 |
| RyhB-TT | 4.36 |
| 17-nt | 8.27 |

### YicC activity on *E. coli* RyhB RNA

A report from the Gottesman lab suggested that YicC was involved in the decay of the iron-responsive, ∼90-nt RNA, RyhB (Chen et al. 2021). We also reported that, in an *E. coli* strain carrying a medium-copy plasmid expressing YicC, the level of RyhB was down ∼10-fold (Ingle et al. 2022). The same effect was observed with overexpressed *B. subtilis* YloC. In addition, a YicC-family member was shown to be involved in degradation of a ∼230-nt RNA in *C. difficile* (Martins et al. 2025). These findings are important as they are currently the sole examples of an *in vivo* function for YicC and related enzymes. Thus, we tested whether RyhB RNA is a substrate for YicC cleavage *in vitro*.

The RyhB RNA folding schematic shown in Fig. 4A is the structure presented in the original RyhB publication (Masse and Gottesman 2002), with 4 U residues added at the 3’ end. The RyhB gene sequence was amplified by PCR from *E. coli* genomic DNA, using an upstream primer that included a T7 RNA polymerase promoter. RyhB RNA was transcribed *in vitro* and incubated with an equimolar amount of YicC enzyme. The reaction products were run on a 6% denaturing polyacrylamide gel and detected by ethidium bromide staining. The result (Fig. 4B) showed that there was activity on this RNA substrate, with two large bands (A and B) and a much smaller band (C) appearing at 2 min of the reaction.

**Fig. 4.**
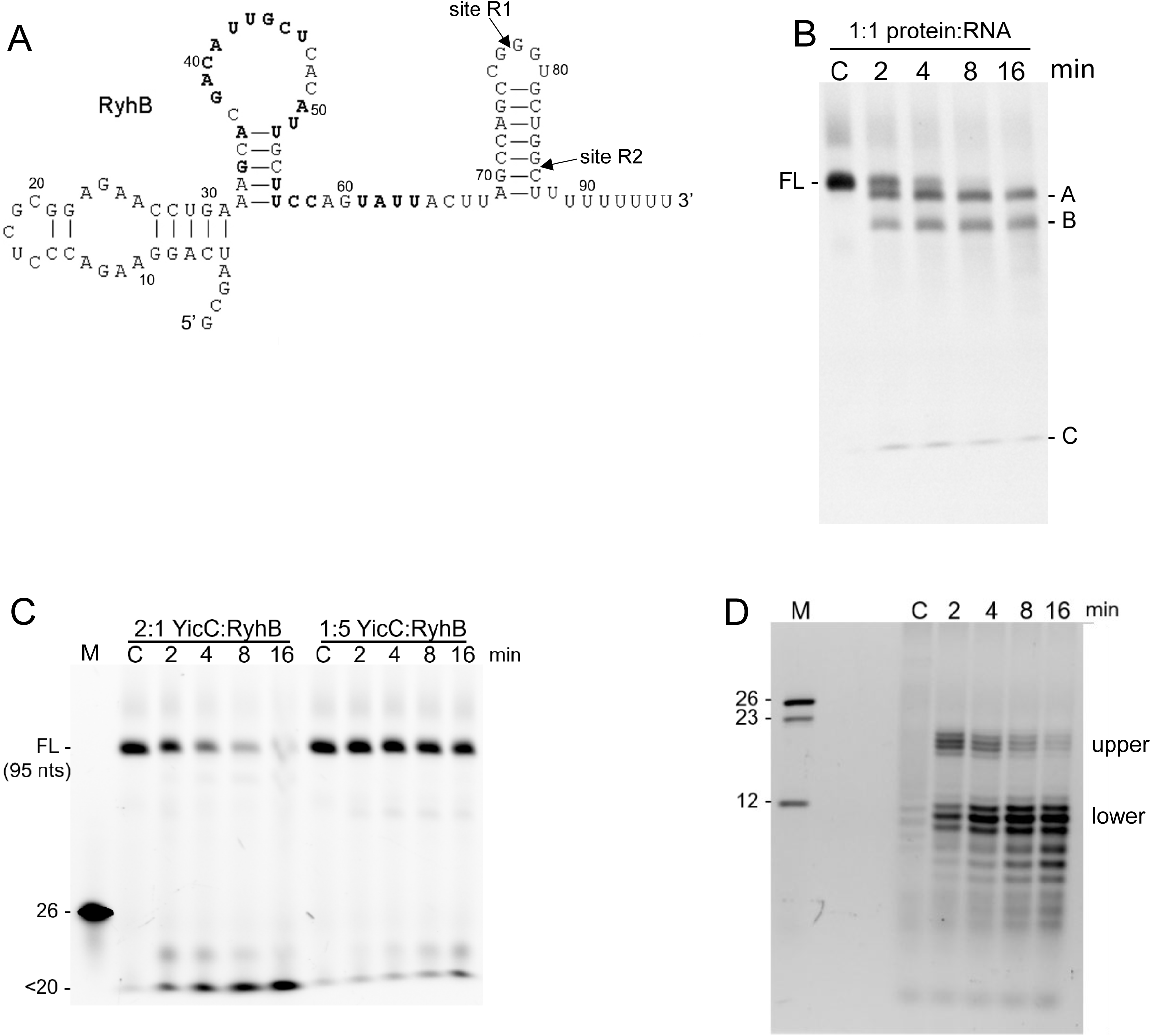
(A) Predicted secondary structure of RyhB RNA. (B) Ethidium bromide-stained gel (6% denaturing polyacrylamide) of products of RyhB RNA incubated with equimolar amount of YicC for the indicated times. FL = full-length RNA. Three cleavage products (A, B, and C) are indicated on the right. (C) YicC activity on 3’-labeled RyhB RNA, using either a 2:1 ratio of YicC:RyhB RNA or a 0.2:1 ratio, run on a 6% denaturing polyacrylamide gel. Marker lane (M) contained 3’-labeled oligo 2051 (26 nts). Sizes of full-length RyhB RNA and the small cleavage product are indicated on the left. (D) Resolution of 3’-labeled band C on a 20% denaturing polyacrylamide gel. Sizes of 3’-labeled oligos in the marker lane (M) indicated at left.

To determine the nature of the RyhB cleavage products, RyhB RNA was labeled at its 3’ end with FTSC. A YicC RNase assay was performed using either a 2:1 molar ratio of YicC:RyhB or 10-fold less YicC (Fig. 4C). For the 2:1 molar ratio reaction, full-length RNA was fully cleaved by 16 min and only the small RNA band, which was smaller than the marker 26-nt oligo, remained. Bands A and B (Fig. 4B), therefore, do not contain the 3’ end of the RNA. Band B is likely the upstream product of cleavage near the 3’ end that gives the 3’-end-labeled small band (band C for the unlabeled RNA), but the nature of band A, which was nearly full-length in size, was not clear (see below). For the 1:5 molar ratio reaction, the full-length RNA was still mostly present after 16 min incubation. This result suggested that a relatively large amount of enzyme was needed to see substantial YicC activity on this substrate, i.e., RyhB RNA is a much weaker substrate than the oligos we have been using (compare with activity on oligo 2051, where a 1:30 YicC:RNA ratio routinely gave substantial cleavage of the RNA substrate after 8 min incubation).

To determine more accurately the nature of RyhB cleavage product C, the YicC reaction products from 3’-labeled RyhB RNA were run on a 20% denaturing gel. The result (Fig. 4D) showed that the “single” band C running at the bottom of the 6% polyacrylamide gels resolved into two sets of bands arising from multiple endonucleolytic cleavages. The upper set of bands (centered around 18 nts) faded with time of the reaction, while the lower set of bands (centered around 10 nts) increased with time of the reaction. Instead of specific bands (as seen in the M lane of Fig. 4C, which is the result of YicC cleavage of oligo 2051), the upper “band” is a cluster of four bands, while the lower “band” is a cluster of bands ranging in size from ∼11 nts to ∼3 nts. The sizes of the observed bands were compatible with cleavages in the predicted 3’ proximal stem-loop which runs from nts 69-87 (Fig. 4A). We hypothesize that cleavage in the loop of this region (Fig. 4A, around site R1) would give 3’ end-labeled fragments of ∼18 nts, while maintaining the stem structure. These 3’-proximal fragments (which separate from the upstream portion of the stem in the denaturing gel but remain paired in the stem during the reaction) decrease in intensity as the reaction continues, likely because they are being cleaved at around site R2. The lower fragments, then, are generated both from cleavage of the full-length RNA and cleavage of the fragments left by cleavage at site R1, and they increase in intensity as the reaction proceeds. Returning to the gel of unlabeled RyhB RNA (Fig. 4B), it appears that band A (which is actually multiple, closely spaced bands) is the result of cleavage at site R2, and band B is the result of cleavage at site R1.

The non-specific nature of YicC cleavage and the need for a high concentration of enzyme to observe cleavage suggested that RyhB RNA binds weakly to the enzyme. This was confirmed by a fluorescence anisotropy competition assay, in which excess RyhB RNA was incubated with a fixed amount of a Cy3-labeled oligo that is a substrate for YicC. While oligo 2051 competed well for binding to YicC protein, no binding of RyhB RNA to YicC was detectable in this assay (data not shown). We conclude that RyhB RNA is a poor substrate for YicC, and the cleavages observed near the 3’ end can be categorized as non-specific cleavages that do not reflect catalytic activity of an enzyme.

A final experiment with RyhB RNA was to test YicC activity on the transcription terminator sequence alone, consisting of nucleotides 66-91 of the sequence shown in Fig. 4A. The size of this RNA, which was designated oligo RyhB-TT (for Transcription Terminator), was 26 nts, the same size as oligo 2051. Initial attempts to label RyhB-TT were unsuccessful, suggesting that the RNA adopts a highly structured conformation that shields the 3’ end and prevents labeling. The predicted stability of RyhB-TT, by Mfold, is -11.5 kcal/mol. To overcome this issue, we heated and slow-cooled oligo RyhB-TT in the presence of an excess of a DNA oligo that was complementary to nts 66-79 of the RyhB sequence, i.e., the ascending half of the strong stem structure. This allowed labeling of the 3’ end. The DNA “splint” was removed by treatment with DNase I and the labeled RyhB-TT RNA was purified and used for a YicC RNase assay. No cleavage of the substrate was observed after 16 min (data not shown). A fluorescence anisotropy assay showed that RyhB-TT oligo was bound YicC several orders of magnitude less strongly than oligo 2051 (Table 1).

### YicC activity on oligos with 5’ and 3’ extensions

Since we observed robust and specific cleavage of various 26-nt RNAs but non-specific cleavage of the 95-nt RyhB RNA, we tested the effect of substrate size by adding nucleotides to oligo 2051 at the 5’ or 3’ end. Oligos 5’+5 and 5’+10 contained the 2051 sequence with five or ten A residues, respectively, added at the 5’ end. The YicC activity assay in Fig. 5A indicated that the addition of even five residues at the 5’ end significantly inhibited the rate of cleavage activity (compare 8 min time points) while not affecting the sites of cleavage. A graph of the data from three independent experiments (Fig. 5B) clearly shows the negative effect of adding nucleotides at the 5’ end.

**Fig. 5.**
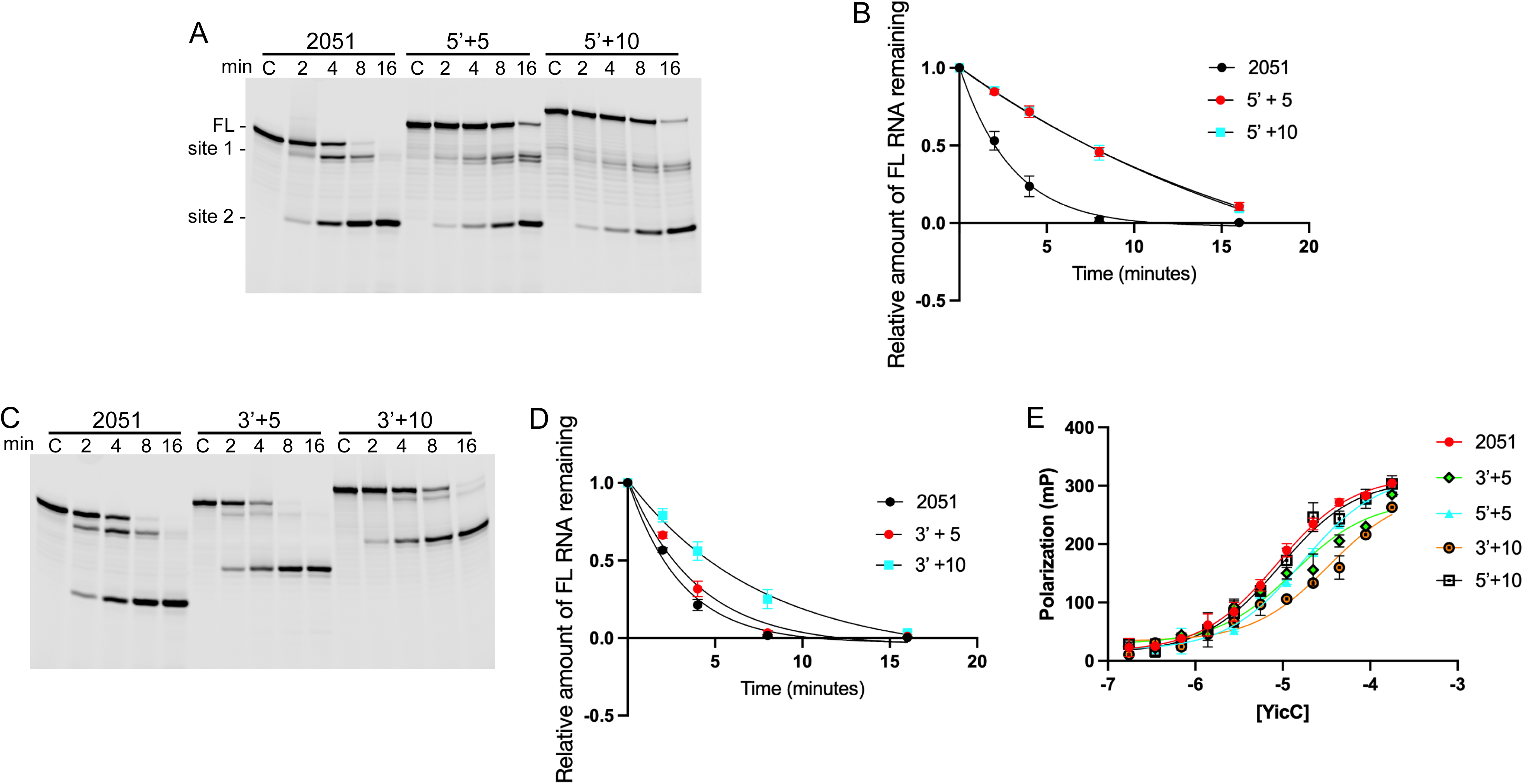
Effect of residues added to the 5’ or 3’ end of oligo 2051. (A) Time course of YicC activity on 3’-end labeled oligos. Oligo 5’+5 had 5 A residues added to the 5’ end; oligo 5’+10 had 10 A residues added to the 5’ end. (B) Decay curves for oligos 5’+5 and 5’+10, relative to oligo 2051. Curves are the average of three experiments. (C) Time course of YicC activity on 3’-end labeled oligos. Oligo 3’+5 had 5 A residues added to the 3’ end; oligo 3’+10 had 10 A residues added to the 3’ end. (D) Decay curves for oligos 3’+5 and 3’+10, relative to oligo 2051. Curves are the average of three experiments. (E) Fluorescence anisotropy measurements on oligos with extensions at the 5’ or 3’ end.

We then examined the effect of adding nucleotides to the 3’ end (Fig. 5C). While the addition of five A’s (oligo 3’+5) had little or no effect (Fig. 5D), the addition of ten A’s (oligo 3’+10) had a measurable effect, although the decline in the decay rate was not as dramatic as when residues were added at the 5’ end. Again, the sites of cleavage were unaffected by the added nucleotides.

To determine whether the effects on YicC activity were due to weaker substrate binding, fluorescence anisotropy experiments were performed using the 3’-end labeled oligos that had been purified by gel extraction. The oligos with a 5-nt or 10-nt extension at the 5’ end had K_D_ values that were ∼1.5-2-fold higher than oligo 2051 (Fig 5E, Table 1). Thus, the significantly slower decay rate of these oligos (Fig. 5B) is likely due not only to an inhibition of the cleavage reaction.

The oligo with a 5-nt extension at the 3’ end had a K_D_ that was ∼1.5-fold higher than oligo 2051, and its decay rate was similar to that of oligo 2051. On the other hand, the oligo with a 10-nt extension at the 3’ end had a K_D_ that was ∼4-fold higher than that of oligo 2051. Thus, the slower decay rate of the 3’+10 oligo is likely due to poor binding.

### YicC activity on a small RNA containing the 5’-proximal hairpin sequence of oligo 2051

Finally, we assayed YicC activity on an RNA that was smaller than 2051 but was predicted to contain the 5’-proximal hairpin structure. For this, we used a 17-nt oligo, which consisted of the first 17 nucleotides of oligo 2051. As shown in Fig. 6, the 17-nt oligo was barely cleaved after 16 min incubation with YicC, indicating that the enzyme’s substrate needed to be a minimum size. To determine the binding affinity, we performed fluorescence anisotropy on the 17-nt oligo vs. oligo 2051. The K_D_ for the 17-nt oligo was more than 8-fold greater than for oligo 2051. Thus, binding of an RNA to YicC appears to require a size greater than 17 nucleotides.

**Fig. 6.**
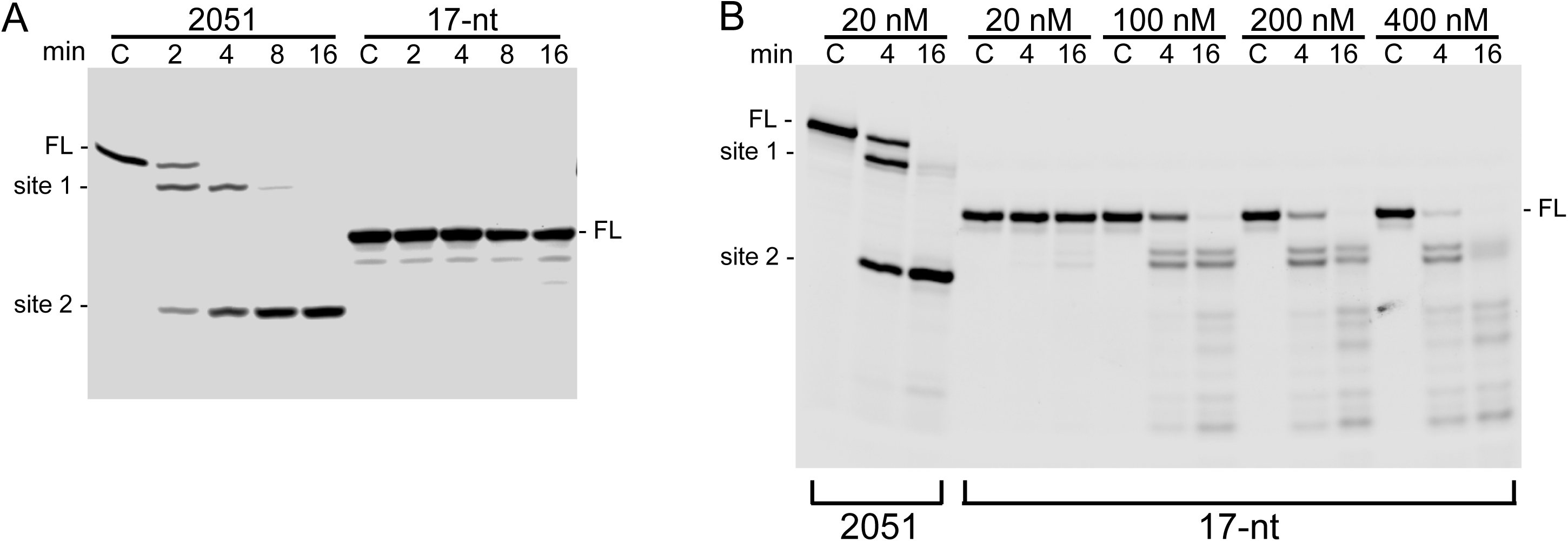
Activity of YicC on a shorter oligo. (A) The 17-nt oligo consisted of the first 17 nucleotides of oligo 2051. Very little cleavage activity was detected under standard assay conditions. (B) RNase assay on the 17-nt oligo with increasing concentrations of YicC.

The 17-nt oligo was used to show directly the effect of assaying YicC activity in the presence of excess protein. Our standard assay contained 20 nM YicC concentration. This gave robust cleavage of oligo 2051 but almost no cleavage of the 17-nt oligo after 16 min incubation (Fig. 6A). Assays for YicC activity on the 17-nt oligo with 5-, 10- and 20-fold more enzyme showed a different pattern (Fig. 6B). With 5-fold more enzyme concentration, cleavage of the substrate at the 4 min time point was observed, as well as many non-specific cleavage products; complete cleavage of the full-length oligo was seen at the 16 min time point. With higher enzyme concentrations, more robust cleavage of full-length RNA and appearance of multiple cleavage products occurred earlier in the time course.

## DISCUSSION

We are at the early stages of understanding the function of YicC-family members and why this enzyme is so highly conserved in eubacteria. We mentioned in the Introduction the possible involvement of YicC in regulation of *E. coli* RyhB RNA and sporulation in *C. diff.* In addition, the gene encoding *E. coli* YicC is located in a set of three contiguous genes that are involved in stationary phase growth and required for growth at high temperature (Poulsen and Jensen 1991). In *B. subtilis*, a Yeast-2-Hybrid screen of protein-protein interactions showed that YloC interacts with YqiB, a subunit of an exodeoxyribonuclease that is likely involved in DNA replication (Marchadier et al. 2011). In the current study, multiple RNA oligos were used to probe aspects of YicC endonuclease activity *in vitro*. We found that YicC cleaves specifically in RNA sequences that are predicted to adopt a double-stranded arrangement. A single nucleotide change in the sequence of the standard oligo 2051 that was predicted to substantially lower the free energy of the 5’-proximal hairpin structure eliminated cleavage at that site (Fig. 2, oligo 3(A)). Furthermore, cleavage in a 5’-proximal hairpin was indifferent to the sequence at that site. We observed specific cleavages in several oligos that had different sequences surrounding cleavage site 1 (cf. Fig. 2). This is unlike the assertion in Huang et al. (Huang et al. 2023) that YicC recognizes specifically a GUG sequence, and prefers a CGUG sequence. We note that for the oligos used in the Huang et al. study, as predicted by Mfold, there was a good correlation between specific cleavage and predicted strength of secondary structure. Overall, 5’-end-labeled RNA oligos that were cleaved by YicC to give a single small product are predicted by Mfold to adopt a structure with lower free energy than RNA oligos that were cleaved non-specifically. Thus, it is possible that changes in the oligo sequence that affected YicC cleavage efficiency in the Huang et al. study were due to an effect on structure rather than the sequence itself.

In this regard, we note that a predicted stability of only -1.8 kcal/mol for the 3’-proximal hairpin (Fig. 1B) as well as the three-nt loop size of that hairpin, which is unfavorable for stem-loop formation (Groebe and Uhlenbeck 1988), make it unlikely that cleavage site 2 of oligo 2051 is recognized as an intramolecular hairpin. Rather, very recent studies in our laboratories indicate that cleavage site 2 is formed when two molecules of oligo 2051 dimerize to form a structure that puts this cleavage site in a double-stranded region (manuscript in preparation). Future work on the characterization of YicC target sites will benefit by inclusion of direct examination of RNA structure, e.g., by NMR, to confirm *in silico* predictions.

Addition of various chemical groups to the 5’ or 3’ end did not affect YicC target site selectivity. The presence of fluorescent dyes of varying molecular weight at either the 5’ end (IRdye 800CWNHS ester, 1166 MW (Ingle et al. 2022), and 5-IAF, 515 MW), or at the 3’ end (FTSC, 458 MW) did not affect where cleavage occurred. We have also visualized YicC cleavage of unlabeled oligo 2051, by staining the gel containing reaction products with SYBR Green, and the same pattern of endonuclease cleavage was observed (data not shown). On the other hand, extending the oligo 2051 sequence at the 5’ end by 5 or 10 A residues had a significant negative effect on YicC RNase activity (Fig. 5A, 5B). Extensions on the 3’ side had less of an effect, but adding 10 A residues made for less efficient cleavage of the full-length RNA (Fig. 4C, 4D). although less than the effect at the 5’ end. Fluorescence anisotropy analysis to measure binding affinity indicated that extensions on the 5’ side (5’+5 and 5’+10) and the 5-nt extension on the 3’ side (3’+5) did not dramatically change the K_D_ (Table 1). Thus, the large decrease in cleavage efficiency of the 5’+10 oligo was likely not due to a deficiency in binding but to an inhibition of the cleavage reaction, possibly by affecting the geometry that places the site 1 cleavage site in the enzyme active site. On the other hand, the 10-nt extension on the 3’ side (3’+10) caused almost a four-fold increase in K_D_, but this was not reflected in a large negative impact on cleavage rate.

While the 5’ and 3’ extensions affected to some extent the rate of cleavage, the site of specific cleavage was maintained in all cases. On the other hand, the much larger RyhB RNA did not bind YicC well, as evidenced by a competition assay, and was cleavage non-specifically and only in the presence of excess enzyme (Fig. 4). Previous reports of accelerated decay of RyhB RNA *in vivo* due to overexpression of YicC or YloC (Chen et al. 2021; Ingle et al. 2022), as well as decay of a ∼230-nt RNA in *C. diff.* due to a YicC homologue (Martins et al. 2021), could be manifestations of an indirect effect (e.g., recruitment of other RNases, as speculated by Chen et al.). Alternatively, *in vivo* there is an additional protein(s) that allows YicC-family members to bind and cleave larger RNAs.

When we analyzed the effect of making targeted changes in the 5’-proximal hairpin (Fig. 3), we observed that increasing the number of base-pairs in the stem from four to six, in oligo 5’stem(6), resulted in a very large (>11-fold) decrease in K_D_, relative to the 5’ stem K_D_ (Fig. 3D; Table 1). This suggests that the affinity of YicC for stem-loop structures increases as the stability of the structure increases. (The predicted folding energy of oligo 5’stem(6) is five logs greater than that of oligo 5’ stem.) On the other hand, little change was observed in the time course of full-length RNA cleavage (Fig. 3B, 3C). Perhaps this can be explained by the fact that, after cleavage of 5’stem(6) at site 1, there are still four base-pairs downstream of the cleavage site, which may delay release of the substrate and decrease overall activity.

When an A residue was introduced into the 5’stem(6) oligo, to give oligo 5’stem(6)+A, the binding affinity was reverted to near that of the 5’stem oligo. However, now we observed that the full-length was cleaved even faster than the 5’stem oligo (Fig. 3B, 3C). There also appeared to be more pronounced minor cleavages in the sequence downstream of the stem-loop structure. We speculate that the preferred substrate for YicC binding is an RNA that has double-stranded structure, but catalysis occurs more efficiently when the structure is able to “breathe,” allowing the scissile bond to be well-situated in the enzyme active site.

We tested one oligo that was smaller than oligo 2051 to approach the question of a minimum size that can be recognized for cleavage. The 17-nt oligo, which consists of the first 17 nucleotides of oligo 251 and is therefore predicted to form the same structure as the 5’-proximal hairpin in 2051, was not a substrate in our standard YicC RNase assay conditions (Fig. 6A). This was also reflected in the >8-fold increase in K_D_ value relative to oligo 2051 (Table 1). We imagine that an RNA that enters the open cage conformation of the YicC hexamer requires a certain size to contact the basic regions that are required for effective binding and subsequent cleavage. Adding excess enzyme, however, overcame these deficits (Fig. 6B). The study of Huang et al. (Huang et al. 2023) used primarily oligos that were 20 nts in length to probe YicC activity, suggesting that this length can also serve as a substrate for YicC. It should be noted, however, that RNase assays in that study used equimolar amounts of RNA substrate and YicC enzyme, and observed only a single 30-min time point, thus making it difficult to compare cleavage efficiencies. At high concentrations *in vitro*, RNases can display different types of cleavage than is observed *in vivo*. For example, *B. subtilis* RNase J1 functions *in vivo* primarily, if not exclusively, as a 5’-to-3’ exonuclease (Condon 2010). However, when there is an excess of RNase J1 present in *in vitro* assays, it acts as a non-specific endonuclease, cleaving at any nucleotide that is in a single-stranded conformation (Daou-Chabo and Condon 2009). The *yloC* gene of *B. subtilis* is expressed at a relatively high level, similar to that of the *pnpA* gene encoding PNPase (Nicolas et al. 2012). Without knowing the biological substrates of YicC-family members, it is difficult to evaluate what sort of enzyme:substrate ratio is needed *in vivo*. Nevertheless, we believe that an *in vitro* condition of a low ratio of enzyme to substrate (e.g., 1:30 in our standard assay) likely reflects the *in vivo* activity of the enzyme. Current efforts are directed at determining the *in vivo* function of the highly conserved family of YicC-like endoribonucleases.

## MATERIALS AND METHODS

### Purification of YicC protein for RNase and anisotropy assays

An overnight culture of the YicC expression strain was diluted 1:400 in 200 ml of LB medium containing 1 mM MgSO_4_, and the culture was grown to OD_600_ of 0.5. YicC expression was induced by addition of IPTG to 400 µM, followed by further incubation with shaking for three hours. Cells were pelleted and stored at -80°C. Cells were grown as described above. Frozen pellets from 100 ml of culture were resuspended in 15 ml ice cold TBS containing 250 mM NaCl. PMSF was added to 1 mM and lysozyme was added to 100 μg/ml. The cell suspension was sonicated (Qsonica) and lysed cells were spun down at 36,000 x g for 30 min. The supernatant was added to 0.3 ml of Ni-NTA and 250 mM imidazole was added for a final concentration of 25 mM imidazole. The lysate and resin were rotated for 1 hr at 4°C. The resin was washed with 20 ml of 50 mM imidazole in TBS/10% glycerol, and protein was eluted with 250 mM imidazole in TBS/10% glycerol. Four buffer exchanges with TBS/10% glycerol were performed in an Amicon-ultra 4 concentrator to a final volume of less than 250 μl. Protein concentration was determined using the Bradford reagent (BioRad), and protein was stored at -80°C. For anisotropy experiments, where more protein was needed, the same protocol was scaled up for 1L cultures.

### RNA 3’ Labeling

RNAs were labeled at the 3’ end essentially as described (Zearfoss and Ryder 2012). 750 pmoles of RNA oligonucleotide (IDT) or *in vitro*-transcribed RNA (HiScribe T7 kit; New England Biolabs) was incubated for 90 min at room temperature in a 50 μl iodination reaction containing 100 mM Na-acetate, pH 5.2, and 100 μM NaIO_4_. NaCl was added to 250 μM, glycogen was added to 0.04 µg/ml, and RNA was precipitated by addition of 2.5 volumes of ice-cold ethanol, incubation at - 20°C for 30 min and centrifugation at 4°C for 25 min at 20,000 x g. A solution of 200 mM FTSC was prepared in dimethyl formamide. RNA was resuspended in labeling buffer containing 100 μM Na-acetate and 6 mM FTSC. The labeling reaction was incubated overnight in the dark at 4°C. The reaction was adjusted to 300 μM Na-acetate, 2.5 volumes of ice-cold ethanol were added and the RNA was centrifuged at 4°C for 25 min at 20,000 x g, washed with 70% ethanol and resuspended in 50 µl of 10 mM Tris, pH 8.0, and 0.1 mM EDTA. Remaining FTSC was removed by clean-up using the RNA Clean & Concentrator-5 kit (Zymo Research).

### RNA 5’ Labeling

RNAs were labeled at the 5’ end essentially as described (Zearfoss and Ryder 2012). 1.5 nmoles of RNA oligo were incubated in a 50 µl T4 polynucleotide kinase (PNK) reaction containing 0.5 mM ATPγS and 5 mM DTT in 1X T4 PNK buffer and 0.4U/µl T4 PNK (New England Biolabs). The reaction was incubated at 37°C overnight. 150 µl of H_2_O was added and the RNA was extracted by addition of equal volume phenol:chloroform:isoamyl alcohol (25:24:1). The mixture was centrifuged for 5 min at 17,900 x *g* and RNA from the aqueous phase was precipitated by addition of one-tenth volume 6M ammonium-acetate, 20 µg of glycogen and 2.5 volumes of cold ethanol. After 30 min incubation at -20°C, RNA was pelleted by centrifugation at 20,000 x *g* for 25 min at 4°C. The pellet was washed with 70% ethanol and resuspended in 42.5 µl of 25 mM HEPES, pH 7.4. To this was added 7.5 µl of 10 mM 5-IAF, prepared in DMSO. Incubation was for 3 hrs at room temperature in the dark, following by purification as described above for 3’ labeling.

### RNase assay

YicC enzyme assays were performed in a 50 μl volume of RNase assay buffer, which was 5 mM MgCl_2_, 50 mM Tris pH 8.0, 7 mM NaCl, 100 mM KCl, and 400 μg/ml BSA. Unlabeled (20 pmoles) and fluorescently labeled RNA (10 pmoles) were used, for a final concentration of 600 nM. YicC protein, containing the N-terminal Sumo tag, was added to a final concentration of 20 nM, giving a 1:30 protein:RNA ratio. At time points after addition of YicC, 10 μl of the reaction were removed into an equal volume Gel Loading Buffer II (Invitrogen) on ice to stop the reaction. Half of each sample (10 μl) was separated on a 20% denaturing polyacrylamide gel (Sequagel; National Diagnostics). Reaction products were visualized on a LI-COR Odyssey F imaging system and analyzed using LI-COR Image Studio software. Decay curves measuring disappearance of full-length RNA over time were graphed using PRISM version 11.0 software.

### Fluorescence Anisotropy

3’-labeled oligos were purified by electrophoresis on a 20% denaturing polyacrylamide gel, staining with ethidium bromide, and cutting out from the gel. Extraction from the gel slice was performed using the ZR small-RNA extraction PAGE Recovery kit (Zymo Research). Labeled oligos were used at a fixed final concentration of 50 nM. YicC protein, which carried the N-terminal Sumo tag, was diluted twofold serially in TBS/10% glycerol, pH 8.0, from 175 µM final concentration across 11 concentrations and a 12^th^ at no protein. After mixing protein in triplicate with RNA on ice, the fluorescent polarization was read on a Victor NIVO plate reader using excitation at 545 nM, emission at 635 nM, and a 595 nM dichroic mirror. The K_D_ values were calculated using Graphpad Prism and a log IC_50_ fit.

## ACKNOWLEDGEMENTS

Support was provided by the National Institutes of Health award numbers GM147211 to D.H.B. and R35GM124838 to M.B.L.

